# Fibroblast-specific genome-scale modelling predicts an imbalance in amino acid metabolism in Refsum disease

**DOI:** 10.1101/776575

**Authors:** Agnieszka B. Wegrzyn, Katharina Herzog, Albert Gerding, Marcel Kwiatkowski, Justina C. Wolters, Amalia M. Dolga, Alida E. M. van Lint, Ronald J. A. Wanders, Hans R. Waterham, Barbara M. Bakker

## Abstract

Refsum disease is an inborn error of metabolism that is characterised by a defect in peroxisomal α-oxidation of the branched-chain fatty acid phytanic acid. The disorder presents with late-onset progressive retinitis pigmentosa and polyneuropathy and can be diagnosed biochemically by elevated levels of phytanic acid in plasma and tissues of patients. To date, no cure exists for Refsum disease, but phytanic acid levels in patients can be reduced by plasmapheresis and a strict diet.

In this study, we reconstructed a fibroblast-specific genome-scale model based on the recently published, FAD-curated model, based on Recon3D reconstruction. We used transcriptomics (available via GEO database with identifier GSE138379), metabolomics, and proteomics data (available via ProteomeXchange with identifier PXD015518), which we obtained from healthy controls and Refsum disease patient fibroblasts incubated with phytol, a precursor of phytanic acid.

Our model correctly represents the metabolism of phytanic acid and displays fibroblast-specific metabolic functions. Using this model, we investigated the metabolic phenotype of Refsum disease at the genome-scale, and we studied the effect of phytanic acid on cell metabolism. We identified 53 metabolites that were predicted to discriminate between Healthy and Refsum disease patients, several of which with a link to amino acid metabolism. Ultimately, these insights in metabolic changes may provide leads for pathophysiology and therapy.

## 1. Introduction

Peroxisomes are organelles that, among other functions, are crucial for cellular lipid metabolism. They perform both anabolic and catabolic processes, including the α- and β-oxidation of very-long-chain fatty acids, dicarboxylic acids, and methyl-branched-chain fatty acids [1]. Furthermore, peroxisomes are involved in the biosynthesis of ether phospholipids, including plasmalogens, bile acids, and essential polyunsaturated fatty acids such as docosahexaenoic acid [2].

Refsum disease is an inborn error of metabolism (IEM) that is caused by biallelic mutations in the gene encoding phytanoyl-CoA 2-hydroxylase (PHYH), resulting in defective α-oxidation of the branched-chain fatty acid phytanic acid (3,7,11,15-tetramethylhexadecanoic acid) [3]. Phytanic acid contains a 3-methyl group and is therefore not a substrate for peroxisomal β-oxidation. Consequently, phytanic acid first needs to undergo α-oxidation, thereby producing pristanic acid, which then can be further degraded by β-oxidation [2]. An alternative metabolic pathway for the breakdown of phytanic acid is ω-oxidation, which takes place in the endoplasmic reticulum [4]. The end product of ω-oxidation of phytanic acid is 3- methyladipic acid (3-MAA), and ω-oxidation has been described to be upregulated in patients with Refsum disease [5]. Refsum disease was first described in 1945 and is clinically characterised by progressive retinitis pigmentosa, polyneuropathy, cerebellar ataxia, and deafness [5]. Biochemically, Refsum disease is diagnosed by elevated levels of phytanic acid in plasma and tissues. Phytanic acid solely derives from the diet, and patients with Refsum disease are mostly diagnosed in late childhood [3,5]. To date, patient management focuses on the reduction of phytanic acid levels by plasmapheresis and a strict diet to reduce the intake of dairy-derived fat [6].

Recently, computational models have become valuable tools to study the complex behaviour of metabolic networks. One type of computational models is genome-scale models of metabolism, which contain all currently known stoichiometric information of metabolic reactions, together with enzyme and pathway annotation [7]. These models can further be constrained and validated by incorporation of different types of data, including mRNA and metabolite profiles, as well as biochemical and phenotypic information [8]. To date, the most comprehensive human models are Recon 3D [9] and HMR 2.0 [10], which are consensus metabolic reconstructions that were built to describe all known metabolic reactions within the human body. Besides, a few tissue- and cell-type-specific models have been developed by incorporating tissue- or cell-specific transcriptomics and proteomics data. These models can be used to predict possible ranges of metabolic fluxes for all enzymes in the network. Flux ranges in diseased and control models can be compared to discover functional changes in the metabolic network. These may be used as biomarkers or give insight into the biochemical origin of disease symptoms [11–15].

In the last decade, a paradigm shift occurred in the field of IEMs. Today, IEMs are no longer viewed according to the “one gene, one disease” paradigm as proposed more than 100 years ago, but recognised to be complex diseases [16]. However, only few studies using systems biology and multi-omics approaches that are widely used for complex diseases have been published for IEMs [8,9,14,17–22].

In this study, we aim to investigate the metabolic phenotype of Refsum disease at the genome-scale, and to study the effect of phytanic acid on cell metabolism. Cultured fibroblasts contain most metabolic functions present in the human body, and biochemical and functional studies in cultured skin fibroblasts are important tools for the diagnosis of patients with a peroxisomal disorder [23]. Therefore, we reconstructed a fibroblast-specific genome-scale model based on fibroblast specific transcriptomics, metabolomics, and proteomics data, and starting from the recently published Recon3D-based model. We obtained these data from healthy controls and Refsum disease patient fibroblasts incubated with phytol, a precursor of phytanic acid. Since flavoproteins play a crucial role in lipid metabolism, we integrated our recently curated set of FAD-related reactions [20]. The resulting model reflects the *in vivo* situation in fibroblasts and demonstrates the physiological effects of a defective α-oxidation. Ultimately, such insights in metabolic changes may provide leads for pathophysiology and therapy.

## 2. Results

### Model curation and generating a fibroblast-specific model

For this study, we used an updated version of the Recon 3D model in which flavoprotein-related metabolism was curated [20]. This addition was essential for this study because many enzymes in fatty acid metabolism are flavoproteins, which carry FAD as a cofactor. Furthermore, a known alternative route for phytanic acid degradation, ω-oxidation, was not accounted for in Recon 3D. In this pathway, phytanic acid is first converted into ω-hydroxy-phytanic acid, followed by oxidation to the corresponding dicarboxylic acid (ω-carboxyphytanic acid; see Fig. 1A). After activation to their CoA-esters, dicarboxylic acids have been shown to enter the peroxisome via ABCD3 [24], and are then degraded via peroxisomal β-oxidation [25]. It is assumed that ω-carboxyphytanic acid follows the same pathway as an unbranched long-chain dicarboxylic acid. The final product of phytanic acid breakdown via ω-oxidation is 3-MAA, which has been identified in urine from patients with Refsum disease [26].

**Figure 1.**
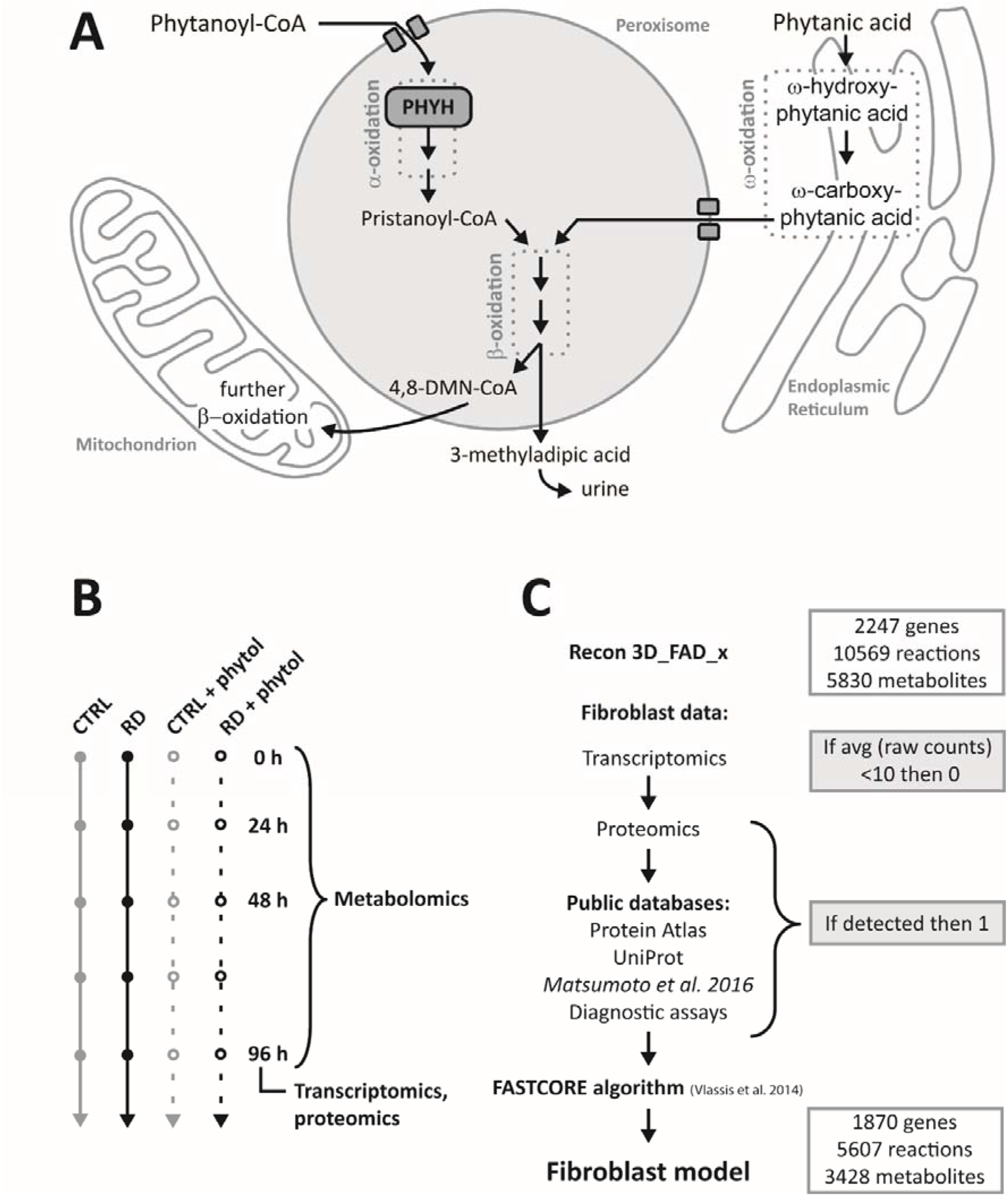
Developing a fibroblast-specific model. A) Schematic overview of relevant metabolic pathways for phytanic acid metabolism. Abbreviations: CoA, coenzyme A; PHYH, phytanoyl-CoA hydroxylase; 4,8-DMN- CoA, 4,8-dimethylnonanoyl-CoA. B) Schematic representation of the experimental setup. Control (CTRL) and Refsum disease (RD) fibroblasts were incubated with or without phytol, the precursor of phytanic acid, for the indicated time points. All cells were seeded and harvested under the same conditions. C) Schematic overview of the steps to obtain a fibroblast-specific model based on constraints of the Recon3D_FAD_x model.

To optimise the model, we added 25 reactions involved in the ω-oxidation and the subsequent β-oxidation of phytanic acid. Furthermore, 17 reactions involved in phytanic acid metabolism were deleted, because they were duplicates of other reactions in the model. Lastly, we examined the import/export reaction boundaries and blocked the flux of several drug metabolism pathways, such as those of statins, ibuprofen, paracetamol, and antibiotics. These pathways were not relevant for this study but could play a role in the model outcome. All changes to the model are summarized in Supplementary Table 1. The resulting curated model was called Recon3D_X_c and is available in on GitHub (https://github.com/WegrzynAB/Papers).

To create a fibroblast-specific model, we generated a fibroblast dataset related to the metabolic genes included in the model. To this end, we cultured human primary control fibroblasts (n=6) and Refsum disease patient-derived fibroblasts with a defect in α-oxidation (n=5) under standardised conditions, and harvested cells after 96 h to isolate RNA and protein. The cells were either incubated with phytol, a precursor of phytanic acid, or with the solvent ethanol (Fig. 1B). Our primary dataset consisted of the data obtained from transcriptomics (RNAseq) and proteomics (shotgun) measurements. In the principal component analysis (PCA), no separation was seen between the groups of fibroblasts (Fig. S1C and D). Differential analysis of the transcriptomics and proteomics data revealed only 12 differentially expressed genes and 18 proteins between the control fibroblasts and fibroblasts defective in α-oxidation (Fig. S1). All differentially expressed genes and 15 proteins were upregulated in the Refsum group relative to controls, while no genes and only three proteins were downregulated. These upregulated genes and proteins were primarily involved in cell cycle control and structure (Supplementary Table 4). When we tested the correlation between protein and RNA levels in the subset of genes that were included in our database, six proteins that were detected in the shot-gun proteomics were not present in the transcriptomics data, even though protein and RNA fractions were obtained from the same sample (Fig. S1C). To complement our own data, we therefore included publicly available information of tissue-specific gene and protein expression levels present in the Human Protein Atlas (Uhlen et al. 2015, www.proteinatlas.org), published transcriptomics and proteomics data obtained from fibroblasts [28], OMIM information [29], fibroblast-specific information published along with the Recon 2 model [8], and information on metabolic assays that are used for diagnostic approaches in fibroblasts (Supplementary Table 2). To generate the fibroblast-specific model, the activity of metabolic reactions was constrained in a two-step manner (Fig. 1C). First, all genes involved in metabolic pathways that were not detected in our transcriptomics data with < 10 raw counts were initially marked as ‘inactive’ (0 in a Boolean vector). Secondly, all these genes were manually cross-examined with our generated database to determine whether the gene was expressed in fibroblasts (either on RNA or protein level). If expressed in skin fibroblasts, the gene was changed to ‘active’ (1 in a Boolean vector). Finally, the FASTCORE algorithm [30] was used to create a flux consistent fibroblast-specific network. This procedure resulted in the final model, ‘fibroblast_CTRL’, which was used for further analysis.

### Model characterisation

First, we tested whether the fibroblast-specific model showed physiological resemblance to fibroblasts *in vivo.* To this end, we used a set of metabolic tasks defined by Thiele et al. [8] and focused explicitly at the metabolic tasks known to be crucial for fibroblast metabolism, i.e. the conversion of glutamine to α-ketoglutarate [31]), or which are known to be absent in fibroblasts, i.e. bile acid metabolism [32]. The fibroblast-specific model completed 208 out of all 419 generic tasks (Supplementary Table 5), demonstrating that the fibroblast model adequately reflects general human metabolism. Additionally, specific reactions known to be present or absent in fibroblasts were also accurately predicted (Table 1), including diagnostically relevant genes (Supplementary Table 5).

**Table 1.** Model performance in the metabolic tasks test. A subset of tasks relevant for fibroblast metabolism selected. For a full list of all tested tasks, see Supplementary Table 5.

| Metabolic task | Active in fibroblasts | Fibroblast |
| --- | --- | --- |
| Bile acid metabolism | NO | NO |
| Pyrimidine degradation | NO | NO |
| Glutamine to citrulline conversion | NO | NO |
| Melatonin synthesis | NO | NO |
| Urea cycle | NO | NO |
| Glutamine conversion to α-ketoglutarate | YES | YES |
| ATP production via electron transport chain | YES | YES |
| Mitochondrial β-oxidation | YES | YES |
| Peroxisomal β-oxidation | YES | YES |
| Peroxisomal α-oxidation | YES | YES |
| ω-oxidation of phytanic acid | YES | YES |
| All 419 generic metabolic tasks |  | 208 |

Subsequently, we simulated the capacity of the fibroblast-specific model to produce ATP from phytanic acid as the single carbon source under aerobic conditions in a minimal medium (consisting of only ions, oxygen, water, and riboflavin). ATP utilisation is explicitly defined in the model and is corrected for ATP investments required for ATP synthesis, such as reactions involved in cofactor synthesis, metabolite transport, and substrate activation. The ATP utilisation flux was used as an objective function of which the value was maximised in the steady-state calculation. Since the flux through the ATP utilisation reaction equals that of ATP production after subtraction of ATP costs at steady state, it reflects the *net* ATP production from a single carbon source (in this case phytanic acid). In contrast to the initial Recon 3D_FAD model, the curated model (Recon3D_FAD_X) and the fibroblast-specific model (fibroblast_CTRL) showed a net ATP production flux of 68.5 and 61.65 mmol · g DW^-1^ · h^-1^, respectively, at a forced phytanic acid uptake flux of 1 mmol · g DW^-1^ · h^-1^.

Furthermore, we created a Refsum disease model (fibroblast_RD) by setting the flux through the phytanoyl-CoA-hydroxylase (PHYH, HGNC:8940) reaction to 0. The fibroblast_RD model was able to metabolise phytanoyl-CoA in minimal medium conditions (Fig. 2, Supplementary Table 3), albeit at a much lower flux than control (38.8 mmol · g DW^-1^ · h^-1^). These results implied that ω-oxidation of phytanic acid and the subsequent β-oxidation in the peroxisomes are less efficient in the ATP production and could require a richer growth media supplemented with glutathione (Figure S2, uptake flux for glutathione was set at 1 mmol · g DW^-1^ · h^-1^). Supplementation of glutathione to the minimum media allowed all studied models to break down phytanic acid, albeit with very strong differences in the total ATP yields. The net ATP production flux of 46.50 mmol · g DW^-1^ · h^-1^ and 86.46 mmol · g DW^-1^ · h^-1^ was shown for the initial Recon 3D_FAD model and the fibroblast_RD model respectively, while much higher net ATP production flux of 116.5 mmol · g DW^-1^ · h^-1^, and 109.3 mmol · g DW^-1^ · h^-1^ was seen for the Recon3D_FAD_X, and the fibroblast_CTRL models (Fig S2). Similarly, we analysed the amino-acid catabolism in the models. All amino acids could be catabolised to yield ATP in the Recon3D_FAD_X model. However, the fibroblast-specific models were unable to metabolise asparagine, histidine, and threonine, as well as nearly no ATP yield from phenylalanine and tyrosine. Furthermore, net ATP production from tryptophan was lower in fibroblast-specific compared to the generic model (Fig. 2). In the minimum media supplemented with glutathione and pantothenic acid all amino acids were broken down; however, asparagine, histidine, phenylalanine, threonine, and tyrosine were showing a strong decrease in the ATP yield in the fibroblast models compared to the generic models (Fig S2).

**Figure 2.**
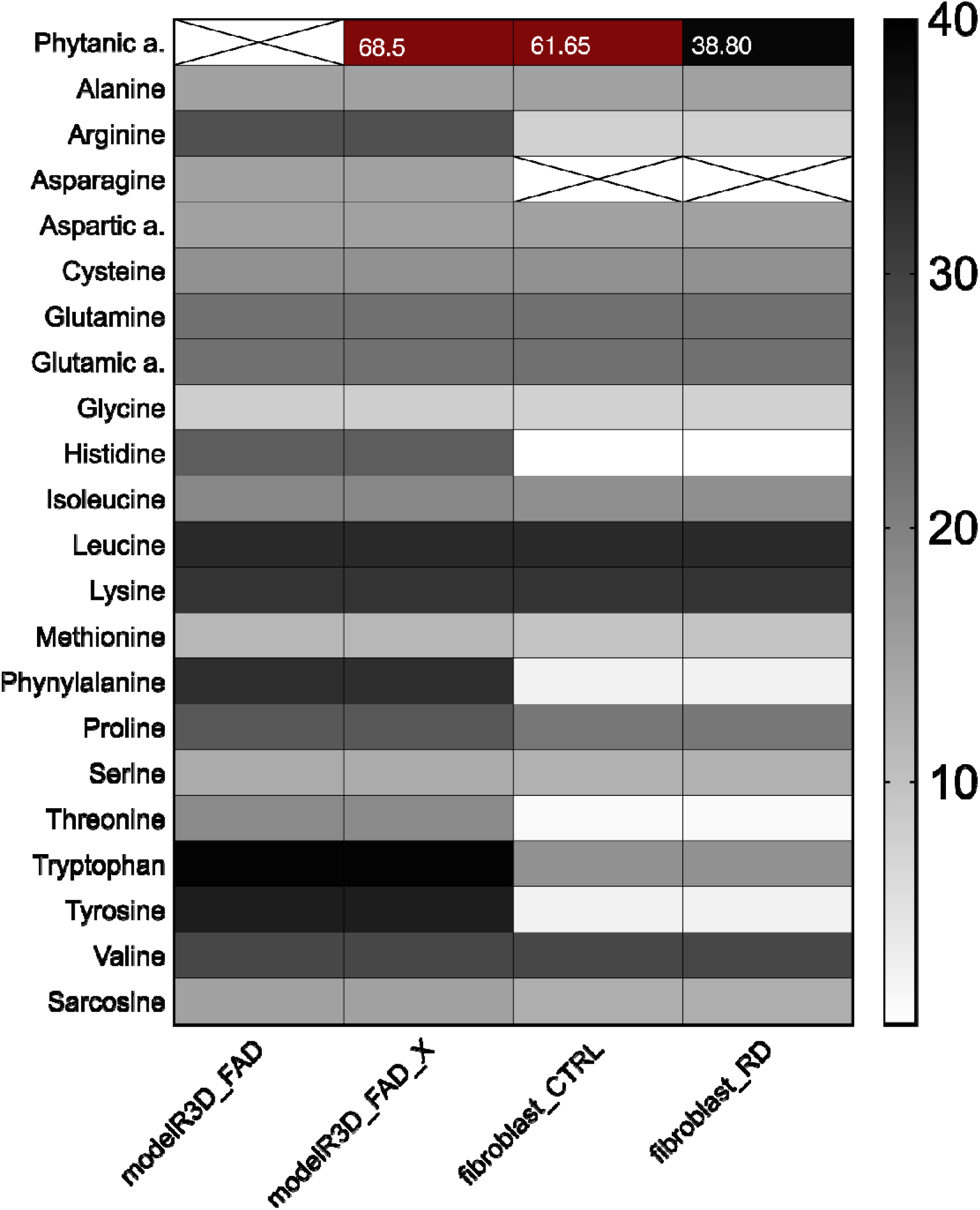
Model predictions of ATP yields from a single carbon source. Assessment of carbon source utilisation on minimal media based on the ATP production from single-carbon source, including Recon3D_FAD, curated Recon3D_FAD for phytanic acid metabolism (Recon3D_FAD_x), the fibroblast-specific model for control (fibroblast_CTRL) and diseased conditions (fibroblast_RD). Grey shades in the heat-maps reflect the relative net ATP production ranging from no (white) to high (black), and very high ATP production (dark red). Crossed-out fields symbolise model inability to metabolise a carbon source (infeasible solution) on minimal media.

To investigate the effect of a defective α-oxidation on the flux distribution in the curated, fibroblast-specific model, we used the fibroblast_RD model to sample the steady-state solution space using the ACHR algorithm [33]. Since genome-scale models typically have multiple steady-state solutions, in this procedure, the solution space reflects the flux ranges found for each reaction when sampling many steady-state solutions (see Methods for details). To be able to compare the results of this analysis with the data from the in vitro fibroblast studies, rich media were used. As expected, the total flux of phytanic acid uptake into the cell was decreased in the fibroblast_RD model when compared to the fibroblast_CTRL model. Because of the simulated deletion of the PHYH gene, α-oxidation was abolished entirely in the fibroblast_RD model, whereas it was active in the fibroblast_CTRL model (Fig. 3). Pathways involved in ω-oxidation, however, were active in both models (Fig. 3). Interestingly, both pathway fluxes were significantly smaller than their maximum rates as obtained from the simulation wherein the maximum flux of an α- or ω-oxidation pathways were used as objective functions (Fig. 3, insert).

**Figure 3.**
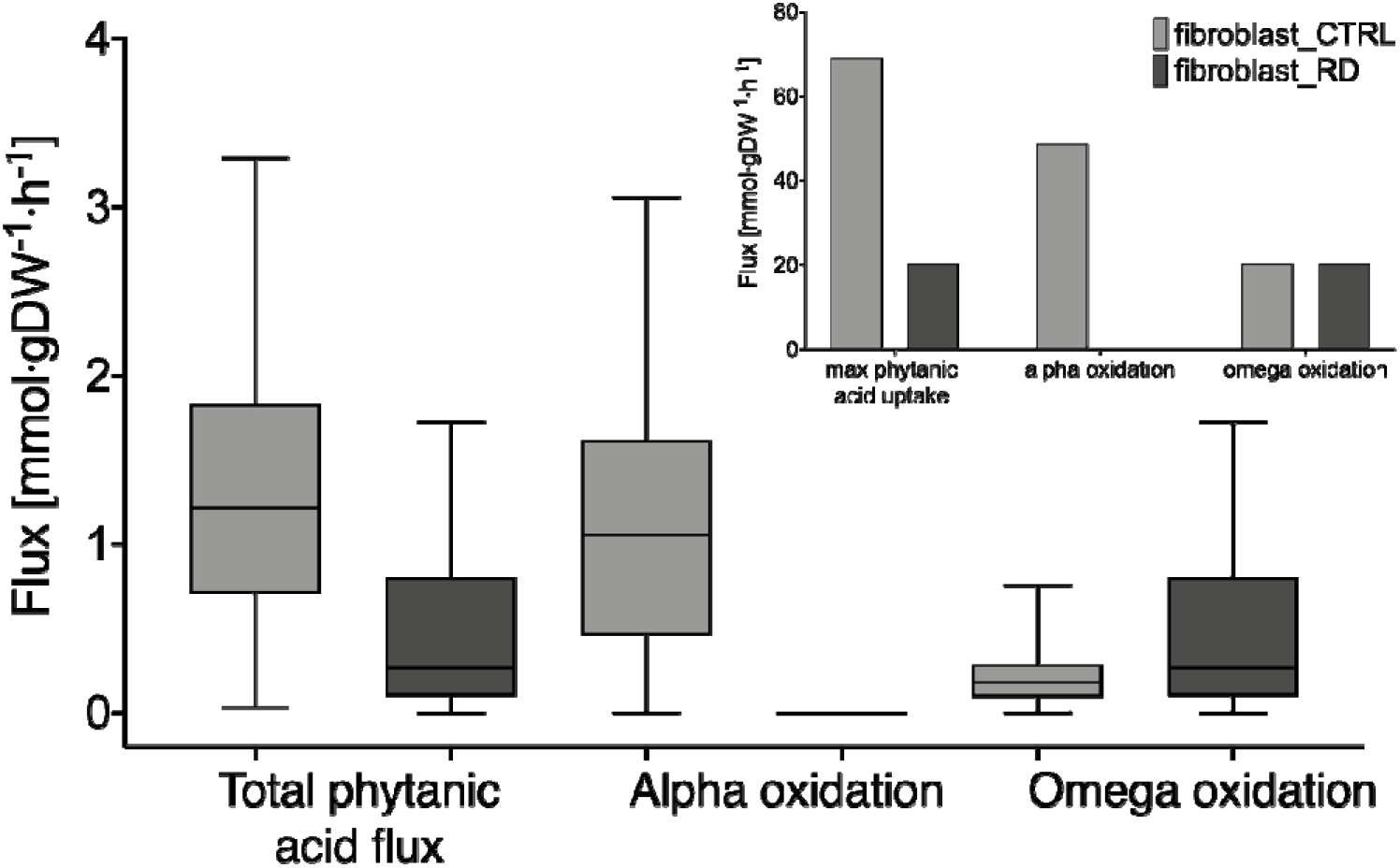
Simulation of phytanic acid metabolism. Missing reactions for phytanic acid metabolism, including α- and ω-oxidation, were added to the Recon3D_FAD model. The curated, fibroblast-specific model shows differences in metabolic fluxes of phytanic acid through the available pathways under normal (control, CTRL) and diseased conditions (Refsum disease, RD). Insert shows maximised fluxes, including blocked α-oxidation in PHYH conditions. Predictions are shown as box-and-whisker (min-max) plots (main figure), or bar plots (insert).

### Metabolic characterisation of fibroblasts cultured *in vitro*

To qualitatively validate our model predictions, we obtained fibroblast-specific metabolomics data. Similar to the transcriptomics and proteomics experiments, we cultured human primary control fibroblasts (n=6) and Refsum disease patient-derived fibroblasts (n=5) under standardised conditions, and collected cell culture medium and cells every 24h for four consecutive days. The cells were incubated with phytol, or with the solvent ethanol (Fig. 1B). First, we measured the levels of total phytanic acid in cells incubated with or without phytol for 96h. The addition of phytol resulted in increased levels of phytanic acid when compared to untreated cells. This was expected, as phytol is converted to phytanic acid once taken up into the cell [34]. In addition, phytanic acid levels were increased in fibroblasts with a defect in α-oxidation when compared to control fibroblasts when phytol was added to the medium (Fig. 2C), reflecting impaired oxidation of phytanic acid.

Furthermore, we measured amino acid profiles in the cell culture medium. We observed no significant changes between the control and Refsum disease groups (Fig. 4B and Fig. S4A) at measured time points. However, a few changes were seen in the rates of uptake or secretion of amino acids (Fig. 4C, and Fig S3). Notably, citrulline and sarcosine have shown to change the directionality in the two groups. While citrulline is secreted, and sarcosine consumed in the healthy fibroblasts exposed to phytol for 96 hours, this situation is reversed in RD fibroblasts. Furthermore, uptake of asparagine is decreased in the RD fibroblasts compared to the healthy ones (Fig. 4C). Other amino acids show some minor differences in their uptake or secretion rates; however, those are not significant (Fig. S3).

**Figure 4.**
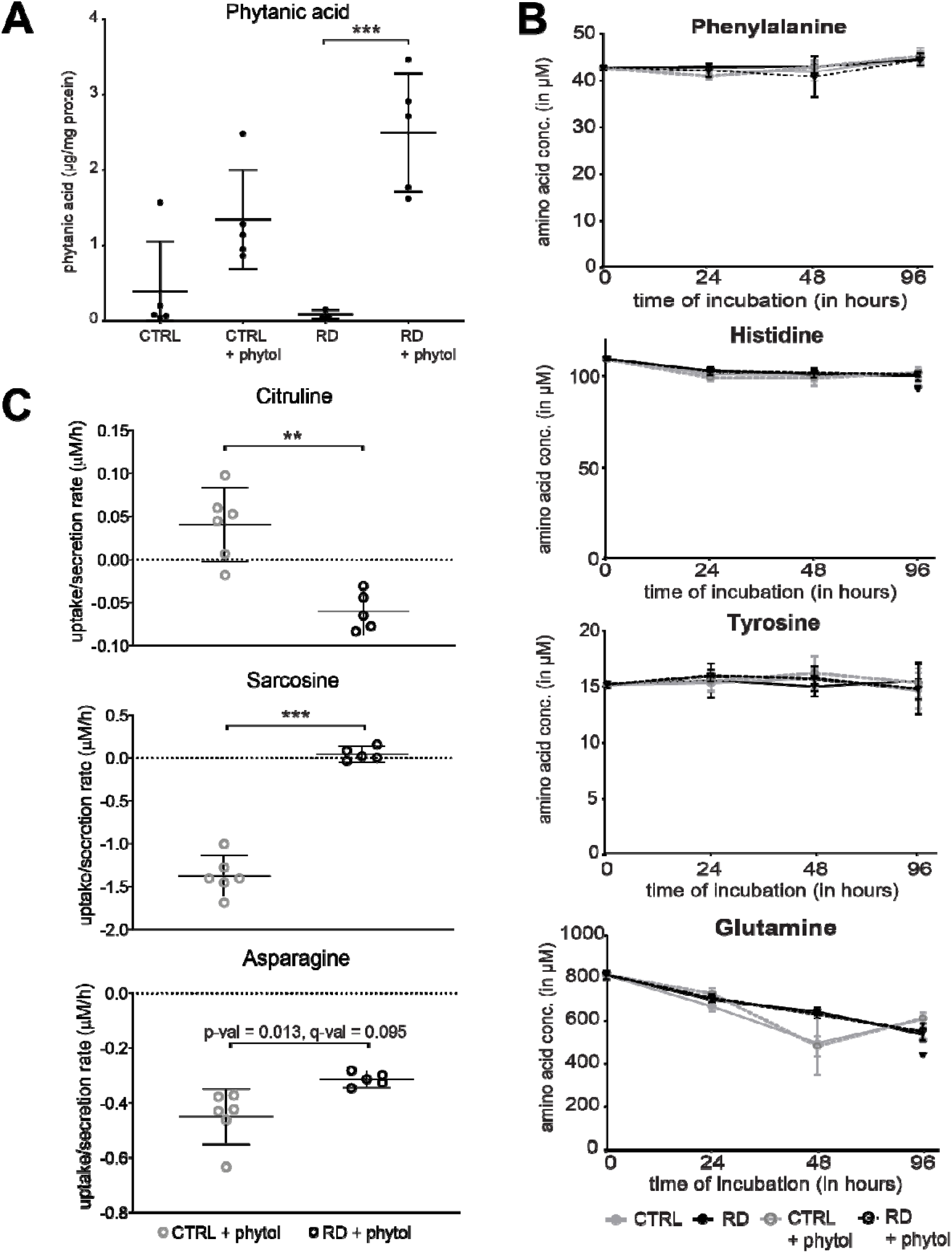
Metabolic characterisation of fibroblasts cultured *in vitro.* Model validation using experimental data of A) phytanic acid, B+C)) amino acid measurements. A) Phytanic acid concentrations were determined in pellets from cultured cells after incubation for 96 hours. Phytanic acid levels are increased in cells incubated with phytol. Per condition, mean per group and 95% confidence interval per group are indicated. Significant differences between the groups were determined by One-way ANOVA (*** p-value < 0.001). B) Significantly changed uptake and secretion rates of amino acids between healthy and RD fibroblasts exposed to phytol for 96 hours. Amino acid concentrations were determined in the medium of the cells 96-hour incubation with phytol. Rates were calculated based on the fresh medium measurements. Significant differences between the groups were determined using a t-test with a two-stage linear step-up procedure of Benjamini, Krieger and Yekutieli, with Q=1%, to correct for the multiple testing (** q-value <0.01, *** q-value < 0.001). Rates of uptake and secretion of other amino acids are shown in Figure S3. C) Amino acid concentrations were determined in the medium of the cells after incubation at indicated time points. Results for other amino acids are shown in Figure S4. D)

Finally, glucose levels (Fig. S2B), cellular protein levels (Fig. S2C), and cell content (Fig. S2D) were similar between the control fibroblasts and the RD fibroblasts with a defect in α-oxidation after 96 hours of cell culture.

### Predicting physiological effects of defective α-oxidation

To investigate other flux changes in the fibroblast_RD model when compared to the fibroblast_CTRL model, we explored the steady-state flux distribution obtained by the sampling of the solution space in the model. We studied changes in the flux ranges of the exchange reactions between control and disease models after forcing a minimum uptake of phytanic acid 0.1 mmol · g DW^-1^ · h^-1^) in the models. Shlomi et al. [19] proposed that if the secretion flux through the exchange reaction is high, it may lead to a high metabolite concentration outside of the cell. In contrast, if uptake is more prevalent, then the extracellular concentration is expected to be lower under the studied conditions. Exchange reactions in the model define the model boundaries. They allow some metabolites to be imported in or secreted from the cell, enabling the model to reach a steady-state. The two models responded differently to the forced phytanic acid uptake flux (Fig 5). The mean value of the phytanic acid flux was reduced by 85% in the fibroblast_RD model when compared to the fibroblast_CTRL model, and secretion of pristanic acid was absent in the RD model (Fig. 5A). The export reaction of 3-MAA, which is the end product of subsequent ω- and β-oxidation of phytanic acid (Fig 4B and Fig 1A), did not show a significant change in its mean flux, while 2,6-dimethylheptanoyl carnitine, one of the end products of canonical degradation pathway of phytanic acid, showed 100% decrease of the flux rate in RD. Besides these known metabolites associated with a defect in α-oxidation, we identified 49 other boundary metabolites that were significantly changed (FDR < 0.05 and log2FC > 1.3) between the fibroblast_RD and the fibroblast_CTRL models (Supplementary Table 6). Of these, 24 flux changes were predicted to lead to higher extracellular concentrations in the absence of PHYH activity, including L-alanine and 3-mercaptolactate-cysteine disulphide (Fig. 5C), caproic acid (Fig. 5D), 2-hydroxybutyrate, and malonyl carnitine, and several di and tripeptides (Supplementary Table 6). On the other hand, 27 distribution flux changes were predicted to result in reduced extracellular concentrations in Refsum disease fibroblasts, such as lactate (Fig. 5C), N-acetyl-asparagine, L-citrulline (Supplementary Table 6) and several di- and tri-peptides (Fig. 5E, and Supplementary Table 6). These changes depend either on the lower/higher uptake rate or on a lower/higher secretion rate (Fig. 5C-E). Interestingly, the rate of secretion of citrulline in our in vitro study showed a significant decrease (Fig. 4C) confirming one of our model predictions.

**Figure 5.**
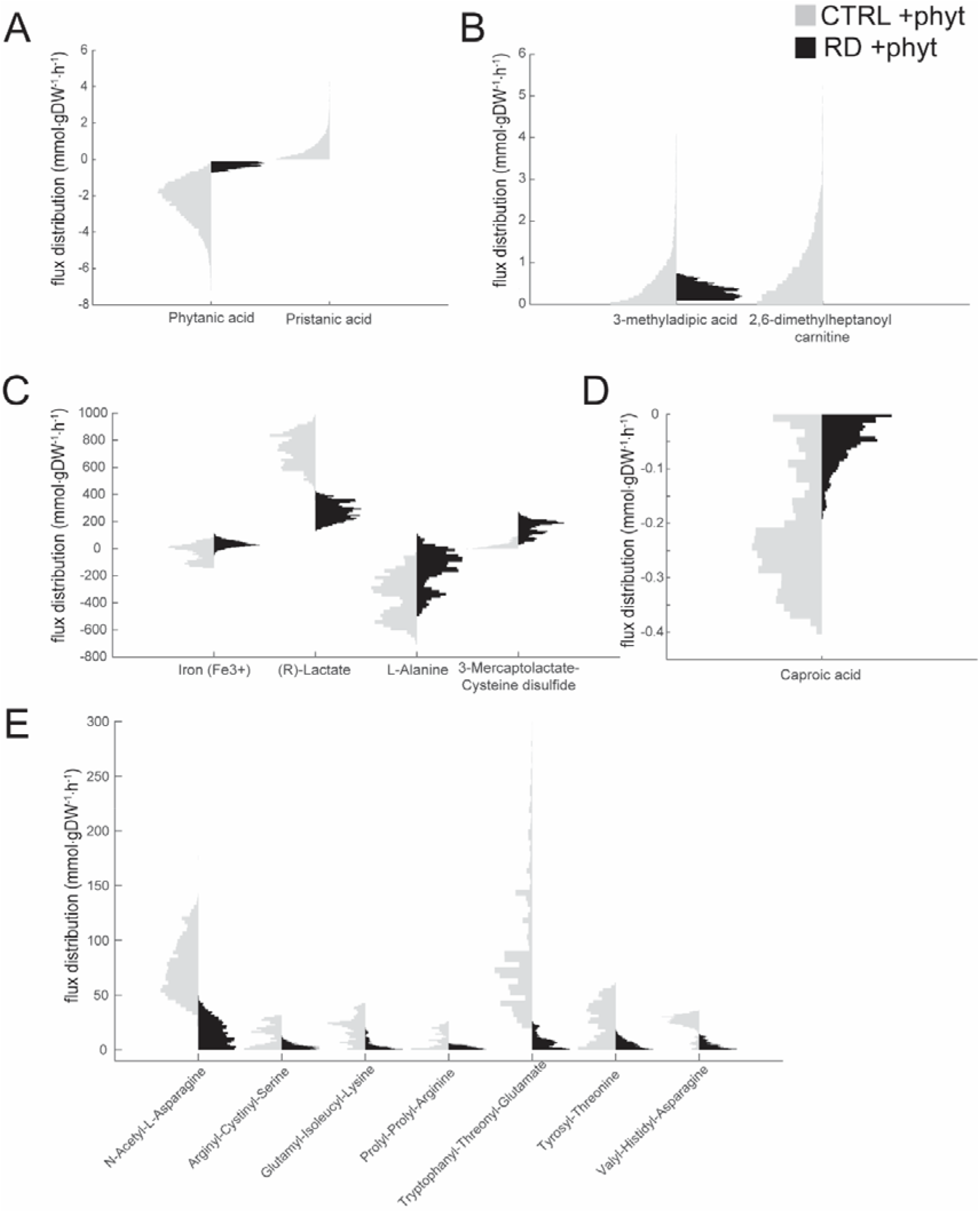
Changes at the level of secretion and uptake reactions between healthy and Refsum models forced to take up phytanic acid. A-E) Secretion /uptake fluxes distributions of metabolites with the most significant differences between Control (CTRL+phyt, grey) and Refsum disease (RD+phyt, black) models forced to take up phytanate selected based on the log2(FC)>1.3 and FDR < 0.05. Statistical differences were analysed using Wilcoxon rank-sum test; FDR values were calculated using Bonferroni-Holm correction.

## 3. Discussion

In this study, we present a fibroblast-specific metabolic model for Refsum disease. Using transcriptomics and proteomics data, we developed a cell-specific metabolic network based on Recon 3D_FAD [20]. Cell-type-specific metabolic models have been reported earlier [13,14,17,21,22,35], and are essential tools to study specific research questions. We studied the effect of phytanic acid loading on the metabolic fluxes in a fibroblast-specific model for Refsum disease, which is characterised by a defect in α-oxidation. Phytanic acid is a natural ligand of peroxisome proliferator receptor α (PPARα) [3,4]. Furthermore, elevated levels of phytanic acid have been reported to induce lipotoxicity in the brain [36]. Many of these findings, however, derive from *in vitro* experiments. The consequences of phytanic acid accumulation have also been studied in a mouse model of Refsum disease, which resembles the clinical symptoms of patients [3,37]. Notably, these mice showed no disease phenotype when fed a regular diet, but only developed the phenotype resembling Refsum disease when challenged with a phytol-enriched diet [37]. Studies in humans, however, are scarce due to limited options for invasive studies. Computational modelling of human cells or tissues is meant to fill this gap partly. In our study, we curated the existing genome-scale model by including pathway information for ω-oxidation and following β-oxidation of phytanic acid and constrained the model to obtain a fibroblast-specific model based on generated data as well as existing databases. The reconstruction of metabolic networks is an iterative process, and updates will assure better accuracy and prediction of the human metabolic model [38].

Using the curated model, we aimed to get an insight into metabolic changes that may provide leads for pathophysiology and biomarkers. Genome-scale metabolic-models have been described to be useful tools for these aims [8–10,14,19–22,35]. In our fibroblast-specific model resembling Refsum disease, the flux of phytanic acid uptake was significantly reduced, reflecting the accumulation of phytanic acid in the body, a known biomarker for Refsum disease [3]. On the other hand, the average 3-MAA secretion rate was not changed between the models. Our results show that it is more desirable for metabolism to lower the phytanic acid uptake rather than increase the ω- oxidation. However, an average sampled flux of 3-MAA secretion was 60 times lower than its maximum theoretical yield (Fig. 3, Fig.5B) showing that the ω-oxidation pathway can be further upregulated, if the uptake of phytanic acid is increased reports that ω-oxidation was upregulated, as described in patients with peroxisomal disorders, complementing partially peroxisomal α- and β-oxidation [4,39].

Besides the changes in the known biomarkers, the model predicted aberrant flux distributions, leading to accumulation or reduction of extracellular metabolites in the Refsum fibroblast model when compared to the healthy model. Interestingly, di and tripeptides were predicted to be changing significantly between the patient and healthy models (Fig. 5F). Biologically active peptides [40] have been found to play important roles in the metabolic functions, including intercellular signal transmission [41] and neuron signal transmission [42,43]. Furthermore, specific peptides are involved in the processes that lead to disease development, and their presence could indicate specific diseases, i.e., serve as disease biomarkers [44–47]. However, the power of the prediction and the value of these metabolic changes in relation to the pathogenesis of phytanic acid in patients with Refsum disease requires further analysis. If validated, our predictions could lead to potential therapeutic strategies to intervene with the accumulation of phytanic acid in these patients. The upregulation of ω-oxidation as an escape route for the breakdown of phytanic acid, and also very-long-chain fatty acids, has been studied in vitro for diseases such as Refsum disease and X-linked adrenoleukodystrophy (ALD) [3,39]. The activation of the cytochrome P450 family (CYP) 4A enzymes, which are known to induce ω-oxidation, has indeed been an attractive target for therapeutic interventions. However, until now, studies using compounds or drugs to upregulate ω-oxidation via CYP4A have not been performed successfully [48]. Our model predicts (Fig. 2 and Fig. S2) that increase in the glutathione levels could not only protect the cells from the oxidative stress postulated to play a role in Refsum disease [49] but potentially also support the phytanic acid breakdown via the ω-oxidation pathway. However, the clinical value of our predictions remains to be evaluated. Fortunately, as mentioned before, a mouse model of Refsum disease exists in which a systemic whole-body effect of phytanic acid accumulation has been studied [37]. Since mice, similarly to humans express ω□hydroxylases from the CYP4 family, the studies of phytanic acid ω-oxidation rates could be performed with either glutathione-enriched diet, or in combination with previously proposed CYP4 inducers: fibrates and statins[4], to determine the clinical potential of our finding.

## 4. Materials and Methods

### Cell culture

Skin fibroblast cell lines from six controls and five patients with genetically confirmed Refsum disease were used. All cell lines were anonymised. Fibroblasts were cultured in 75-cm^2^ flasks for transcriptomics and proteomics analysis, and in 25-cm^2^ flasks for metabolomics experiments. Cells were cultured in Ham’s F-10 medium with L-glutamine, supplemented with 10% foetal calf serum (Invitrogen, Carlsbad, CA, USA), 25 mM Hepes, 100 U/mL penicillin and 100 μg/mL streptomycin and 250 μg/mL amphotericin in a humidified atmosphere of 5% CO2 at 37 □C. Cells were seeded on the same day and incubated for the indicated time points (Fig. 1B). Cells were incubated with 25 uM phytol, dissolved in ethanol, or ethanol as the vehicle. Cells were harvested by trypsinisation (0.5% trypsin-EDTA, Invitrogen) and washed once with phosphate-buffered saline and twice with 0.9% NaCl, followed by centrifugation at 4 □C (16100 x g for 5 min) to obtain cell pellets. For metabolomics experiments, the cell culture medium was collected before harvesting. Cell pellets and medium samples were stored at −80 □C until analysis.

### RNA & protein Isolation for RNAseq and Shotgun proteomics measurements

RNA and protein were isolated from the cell pellets from the T75 cultures using TRIzol™ Reagent (ThermoFisher Scientific) using supplier protocol for RNA and protein extraction. RNA pellets were dissolved in 50uL of RNase free water, and RNA concentrations were measured using NanoDrop™ 2000 Spectrophotometer (ThermoFisher Scientific). Protein pellets were dissolved in 200 uL 5% SDS solution and protein concentrations were determined using Pierce™ BCA Protein Assay Kit (ThermoFisher Scientific).

### RNAseq

#### Sample Preparation and sequencing

First quality check of and RNA quantification of the samples was performed by capillary electrophoresis using the LabChip GX (Perkin Elmer). Non-degraded RNA-samples were selected for subsequent sequencing analysis. Sequence libraries were generated using the Nextflex Rapid Illumina Directional RNA-Seq Library Prep Kit (Bioo Scientific) using the Sciclone NGS Liquid Handler (Perkin Elmer). The obtained cDNA fragment libraries were sequenced on an Illumina Nextseq500 using default parameters (single read 1×75bp) in pools of multiple samples, producing on average 4 million reads per sample.

#### Gene expression quantification

The trimmed fastQ files were aligned to build human_g1k_v37 Ensemble [50] release 75 reference genome using hisat/0.1.5-beta-goolf-1.7.20 [51] with default settings. Before gene quantification, SAMtools/1.2-goolf-1.7.20 [52] was used to sort the aligned reads. The gene-level quantification was performed by HTSeq-count: HTSeq/0.6.1p1 [53] using --mode=union, with Ensembl release 75 [50] was used as a gene annotation database.

#### Calculate QC metrics on raw and aligned data

Quality control (QC) metrics are calculated for the raw sequencing data. This is done using the tool FastQC (FastQC/0.11.3-Java-1.7.0_80) [54]. QC metrics are calculated for the aligned reads using Picard-tools (picard/1.130-Java-1.7.0_80) [55] CollectRnaSeqMetrics, MarkDuplicates, CollectInsertSize-Metrics and SAMtools/1.2-goolf-1.7.20 flagstat.

### Shotgun proteomics

#### In-gel digestion and strong cation-exchange (SCX) fractionation

Protein samples were mixed with LDS loading buffer (NuPAGE) at a concentration of 3.4 μg total protein. The sample was run briefly into a precast 4-12% Bis-Tris gels (Novex, ran for maximally 5 min at 100 V). The gel was stained with Biosafe Coomassie G-250 stain (Biorad), and after destaining with milliQ-H_2_O, the band containing all proteins was excised from the gel. The gel band was sliced into small pieces, washed subsequently with 30% and 50% v/v acetonitrile in 100 mM ammonium bicarbonate (dissolved in milliQ-H_2_O), each incubated at RT for 30 min while mixing (500 rpm) and lastly with 100% acetonitrile for 5 min, before drying the gel pieces in an oven at 37 °C. The proteins were reduced with 20 μL ten mM dithiothreitol (in 100 mM ammonium bicarbonate dissolved in milliQ-H_2_O, 30 min, 55 °C) and alkylated with 20 μL 55 mM iodoacetamide (in 100 mM ammonium bicarbonate dissolved in milliQ-H_2_O, 30 min, in the dark at RT). The gel pieces were washed with 50% v/v acetonitrile in 100 mM ammonium bicarbonate (dissolved in milliQ-H_2_O) for 30 min while mixing (500 rpm) and dried in an oven at 37 °C) before overnight digestion with 20 μL trypsin (1:100 g/g, sequencing grade modified trypsin V5111, Promega) at 37 °C. The next day, the residual liquid was collected before elution of the proteins from the gel pieces with 20 μL 75% v/v acetonitrile plus 5% v/v formic acid (incubation 20 min at RT, mixing 500 rpm). The elution fraction was combined with the residual liquid and was dried under vacuum and resuspended in 30 μL of 20% v/v acetonitrile plus 0.4% v/v formic acid (dissolved in milliQ-H_2_O) for SCX fractionation. Samples were loaded onto an SCX StageTips (20 μL tip StageTip, Thermo Scientific) according to the manufacturer’s instructions, except that the elution solvent (500 mM ammonium acetate in 20% v/v acetonitrile, dissolved in milliQ-H_2_O) plus 0.4% v/v formic acid was used instead of the 1M NaCl solution in this protocol during initialization. After loading and washing of the peptides according to the protocol, the peptides were eluted in three separate fractions by stepwise elutions (30 μL each) of 25 mM, 150 mM and 500 mM ammonium acetate in 20% v/v acetonitrile (dissolved in milliQ-H_2_O). The collected flowthrough was polled with the last elution fraction. The elution fractions were dried under vacuum and resuspended in 8 μL 0.1% v/v formic acid (dissolved in milliQ-H_2_O).

#### LC-MS analysis

Discovery mass spectrometric analyses were performed on a quadrupole orbitrap mass spectrometer equipped with a nano-electrospray ion source (Orbitrap Q Exactive Plus, Thermo Scientific). Chromatographic separation of the peptides was performed by liquid chromatography (LC) on a nano-HPLC system (Ultimate 3000, Dionex) using a nano-LC column (Acclaim PepMapC100 C18, 75 μm x 50 cm, 2 μm, 100 Å, Dionex, buffer A: 0.1% v/v formic acid, dissolved in milliQ-H_2_O, buffer B: 0.1% v/v formic acid, dissolved in acetonitrile). In general, 6 μL was injected using the μL-pickup method with buffer A as a transport liquid from a cooled autosampler (5 °C) and loaded onto a trap column (μPrecolumn cartridge, Acclaim PepMap100 C18, 5 μm, 100 Å, 300 μmx5 mm, Dionex). Peptides were separated on the nano-LC column using a linear gradient from 2-40% buffer B in 117 min at a flow rate of 200 nL/min. The mass spectrometer was operated in positive ion mode and data-dependent acquisition mode (DDA) using a top-10 method. MS spectra were acquired at a resolution of 70.000 at m/z 200 over a scan range of 300 to 1650 m/z with an AGC target of 3e^6^ ions and a maximum injection time of 50 ms. Peptide fragmentation was performed with higher energy collision dissociation (HCD) using normalised collision energy (NCE) of 27. The intensity threshold for ions selection was set at 2.0 e^4^ with a charge exclusion of 1≤ and ≥7. The MS/MS spectra were acquired at a resolution of 17.500 at m/z 200, an AGC target of 1e^5^ ions and a maximum injection time of 50 ms and the isolation window set to 1.6 m/z

#### LC-MS data analysis

LC-MS raw data were processed with MaxQuant (version 1.5.2.8) [56]. Peptide and protein identification was carried out with Andromeda against a human SwissProt database (www.uniprot.org, downloaded November 10, 2016, 20,161 entries) and a contaminant database (298 entries). The searches were performed using the following parameters: precursor mass tolerance was set to 10 ppm, and fragment mass tolerance was set to 20 ppm. For peptide identification two miss cleavages were allowed, a carbamidomethylation on cysteine residues as a static modification and oxidation of methionine residues as a variable modification. Peptides and proteins were identified with an FDR of 1%. For protein identification, at least one unique peptide had to be detected, and the match between runs option was enabled. Proteins were quantified with the MaxLFQ algorithm [57] considering unique peptides and a minimum ratio count of one. Results were exported as tab-separated *.txt for further data analysis.

### Differential analysis of transcriptomics and proteomics

Differential gene/protein expression analysis based on the negative binomial distribution was performed using DESeq2 [58]. Genes for which summed across all samples raw counts were higher than 20 were analysed. Protein intensities were transformed to integers and analysed similarly to the transcriptomics data.

### Cell growth

Fibroblasts were seeded in 96-well plate with a density of 2000cells/well and cultured in 200uL of medium for seven days. xCELLigence system (ACEA Biosciences Inc.) was used to monitor cells attachment and growth in real-time [59]. Areas under the curve were calculated using Prism7 (GraphPad Software).

### Metabolomics

#### Determination of protein concentration in cell pellets

Cell pellets were sonicated in 250uL of water. Protein concentration was determined using the Pierce™ BCA Protein Assay Kit (ThermoFisher Scientific).

#### Amino-acid profile

To analyse the amino-acid profile of medium from cell cultures 100uL of the medium sample was mixed with 100uL of internal standard (12mg of norleucine mixed with 15g sulphosalicylic acid in 250ml of water). The analysis was performed according to the method of Moore, Spackman, and Stein [60] on a Biochrom 30™ Amino acid Analyser (Biochrom.co.uk). Acquisition and data handling were done with Thermo Scientific™ Chromeleon™ 7.2 Chromatography Data System software (ThermoFisher Scientific).

#### Sugar measurements

To analyse sugar profiles, 250ul of the medium sample or 100ul of a standard mix (50mg of D-(+)-glucose in 50ml of water) was mixed with 100ul of internal standard (50mg phenyl-b-D-glucopyranoside in 50ml of water mixed with 1 ml of chloroform). Glucose analysis was performed as described by Jansen et al. [61] on a Trace GCMS (Thermo Fisher Scientific). Acquisition and integrations were done with Xcalibur™software (ThermoFisher Scientific).

#### Phytanic acid measurement

Phytanic acid levels were measured as described previously [62].

### Model curation

Our model is based on a previously published FAD-curated version of Recon 3D [20]. Current representation of phytanic acid metabolism was analysed and compared with current knowledge [4,63]. Missing reactions in omega-oxidation of phytanic acid and follow-up peroxisomal beta-oxidation of its products were added to the reconstruction. Additionally, invalid or duplicated reactions (created by merge of different metabolic reconstructions to create Recon 2 model [8]) were removed. The curated model was saved as Recon3D_FAD_X. For detailed information on all the changes to the model, see Supplementary Table 1 [fix, del].

### Model constrains

We examined all exchange/demand reactions to determine the model constraints. Since drug metabolism introduced by Sahoo et al. [64] is out of the scope of our research, we decided to block the import/export reactions for drugs and their metabolites. Additionally, we identified redundant demand and sink reactions that duplicate some exchange/demand reactions or allow sink reaction for a metabolite whose metabolism has been fully reconstructed and does not create a dead-end pathway. Last, we closed all import reactions besides those that transported compounds present in the culture media, water, and oxygen. All the changes can be examined in the Supplementary Table 1 [constraints].

Additionally, ‘biomass_reaction’ minimum flux was set to 0.1 mmol·gDW^-1^·h^-1^, to mimic the essential cell maintenance (protein synthesis, DNA and RNA synthesis etc.), unless stated otherwise, as in [65]. Other constraints used only in specific simulations are indicated where applicable.

### Fibroblast-specific gene database

A database containing information about the expression levels of metabolic genes (genes present in the metabolic reconstruction Recon 3D_FAD) and proteins in human fibroblasts was first generated based on the results from our transcriptomics and proteomics experiments. Additionally, we added information present in the Human Protein Atlas [27,66], OMIM [29] fibroblast-specific information published along with the first Recon 2 model [8], and UniProt [67] databases. Experimental data from human fibroblast gene expression levels by Matsumoto et al. [28] was also included. Usage of fibroblasts in diagnostics of specific gene defects was also examined. In the end, a binary decision was made about fibroblast-specific genes – 1 if there was evidence for a gene/protein to be present in human fibroblasts, 0 for genes classified as inactive in fibroblasts. Database, including the final decision, is available as a Supplementary Table 2.

### Fibroblast-specific model generation

A list of reactions depending on the genes marked as active was used as a core set for the FASTCORE algorithm [30] implemented in The COBRA Toolbox v3.0 [68]. Next, reactions dependent on the inactive genes were removed, and fastcc algorithm [30,68] was used to generate a flux, consistent fibroblast-specific model. The final model, named ‘fibroblast_CTRL’ is available in our Github folder.

## Model analysis

### Refsum simulations

Phytanoyl-CoA hydroxylase deficiency (Refsum disease) was simulated as a single gene deletion (PHYH, HGNC:8940). Additionally, ω-oxidation (‘CYP450phyt’ reaction) and α-oxidation (‘PHYHx’, reaction) pathways maximum rates were constrained to 20.2176 and 48.7656 mmol·gDW^-1^·h^-1^ respectively, to reflect those described in the literature [69,70]. Lastly, the ‘EX_phyt(e)’ reaction upper boundary was set to −0.1 mmol·gDW^-1^·h^-1^ to force the model to utilise phytanic acid at a minimum rate of 0.1 mmol·gDW^-1^·h^-1^ for the simulations resembling fibroblasts with phytol added to the medium.

To sample the solution space of generated models, ACHR algorithm [33] implemented in the COBRA Toolbox 3.0 [68] was used. Randomly selected 10000 sampled points were saved with from the total of 50000 sampled points with a 500 step size.

### Calculation of maximum ATP yield per carbon source

To calculate the maximum ATP yield per carbon source, we adapted the method developed by Swainston et al. [38]. Shortly, all uptake rates of nutrients were set to 0, except for a set of reactions defined collectively as a minimal medium (Ca^2+^, Cl^-^, Fe^2+^, Fe^3+^, H^+^, H_2_O, K^+^, Na^+^, NH_4_ SO_4_^2-^, Pi, Riboflavin) for which the uptake/export fluxes rates were set to −1000 and 1000 mmol·gDW^-1^·hr^-1^ respectively. For each of the specified carbon sources, the uptake flux was set to −1 mmol·gDW^-1^·hr^-1^ forcing the model to consume it at a fixed rate. The demand reaction for ATP, ‘DM_atp_c_’ was used as an objective function flux, which should be maximised in the optimisation process. The oxygen intake flux was set to ‘EX_o2(e)’ −1000 mmol·gDW^-1^·hr^-1^ to maintain aerobic conditions. If the model was unable to breakdown specified carbon source to ATP, the steady-state flux could not be reached (infeasible solution).

### Statistical analysis of model predictions

Flux distribution of each exchange reaction was compared between the control (CTRL) and Refsum’s (RD) to find the most changed metabolite fluxes. To this end, we tested normality and variance of the distributions using Single sample Kolmogorov-Smirnov goodness-of-fit hypothesis test and Two-sample F test for equal variances, respectively. Depending on the outcome Student’s t-test (for normally distributed samples with equal or unequal variance) or Wilcoxon ranks sum test (for non-normally distributed samples with unequal variance) were used to determine whether the differences between the control (CTRL) and Refsum’s (RD) models were significant. Bonferroni-Holm correction for multiple comparisons was used to calculate the adjusted p-values (FDR). Significance thresholds were set at FDR < 0.05 and log_2_(FC) > 1.3.

## Software

Model curation and all simulations were carried out with MatLab R2019a (MathWorks Inc., Natick, MA) using the Gurobi8.1 (Gurobi Optimization Inc., Houston TX) linear programming solver and the COBRA 3.0 toolbox [68].

## Data availability

The mass spectrometry proteomics data have been deposited to the ProteomeXchange Consortium via the PRIDE [71] partner repository with the dataset identifier PXD015518.

The RNAseq data have been deposited to the GEO database [72] with the identifier GSE138379.

## Author Contribution

ABW, KH, RW, HW, and BMB conceived the idea. ABW and KH designed the experiments. KH, AEML, and ABW performed cell experiments. KH performed the phytanic acid measurement experiment. AG performed the metabolomics experiment. MK and JCW performed the proteomics experiments. AMD, and ABW performed the cell growth experiments. ABW performed the computational modelling. ABW and KH analyzed the data. ABW and KH wrote the first draft of the manuscript. All authors participated in the reviewing and editing of the subsequent drafts.

## Acknowledgements

The authors would like to thank Sarah Stolle for upload of transcriptomics data to the GEO database and, together with Anne-Claire M.F. Martines, and Dirk-Jan Reijngoud, for valuable comments to this manuscript. Additionally, we would like to thank François Bergey for his assistance with differential gene analysis.

This work was supported by the Marie Curie Initial Training Networks (ITN) action PerFuMe [project number 316723] and the University Medical Centre Groningen.

---

**Figure S1.**
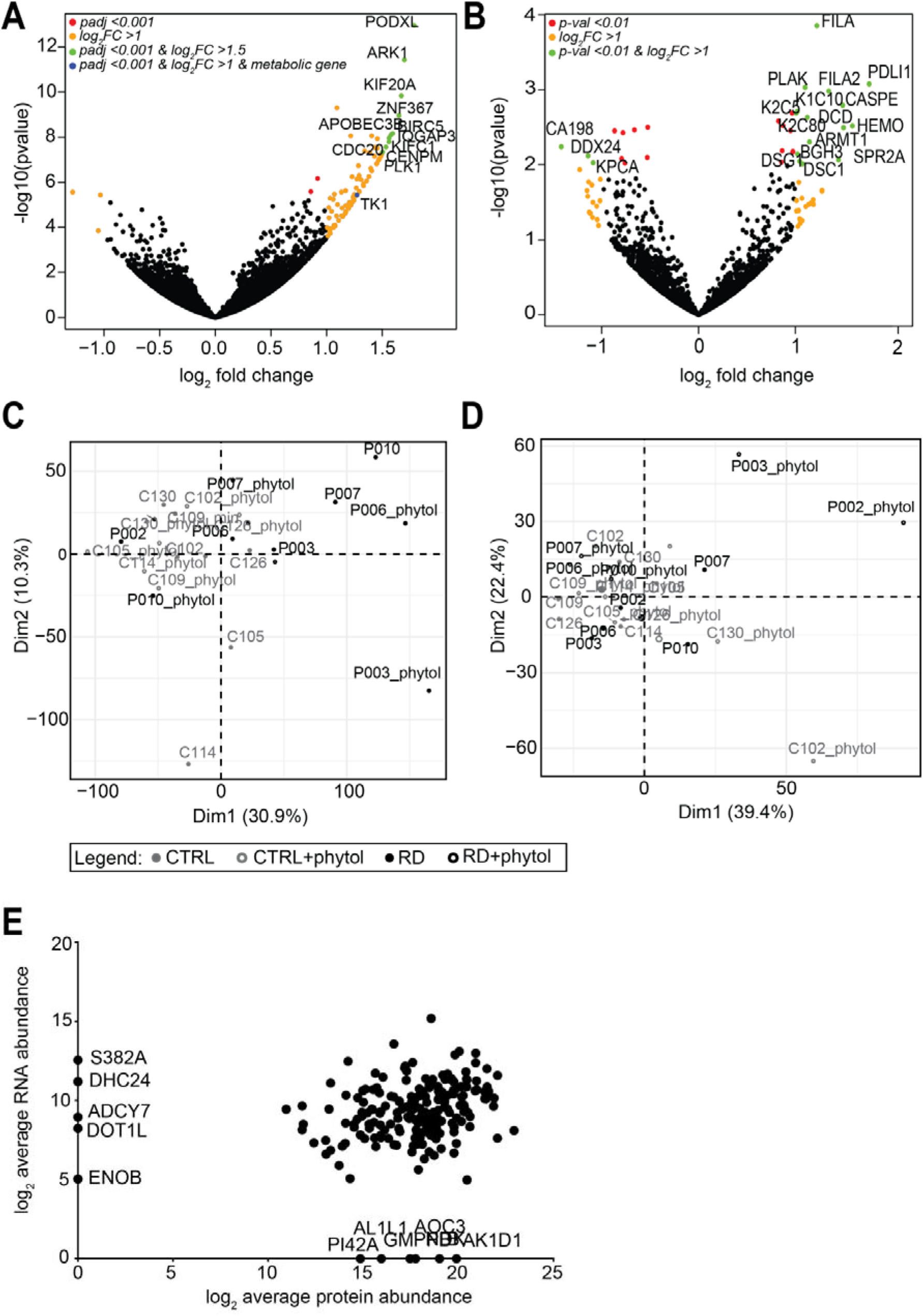
A+B) Volcano plots depicting A) the transcriptomics data, and B) proteomics data derived from fibroblasts incubated with phytol for 96 hours. Genes and proteins, resp., with significant differences in expression between the diseases (Refsum disease) and control groups are indicated with coloured dots. Gene names are shown for genes and proteins, resp., indicated with green dots. Blue dots represent metabolic genes, as included in the Recon3D model, that were expressed differentially at the significance level below 0.001, and their expression levels were changed by minimum 1-fold. C+D) Principal Component Analysis for C) transcriptomics data, and D) proteomics data. E) Correlation plot showing log 2 average abundance of all proteins (x-axis) and genes (y-axis) that were included in the Recon 3D model.

**Figure S2.**
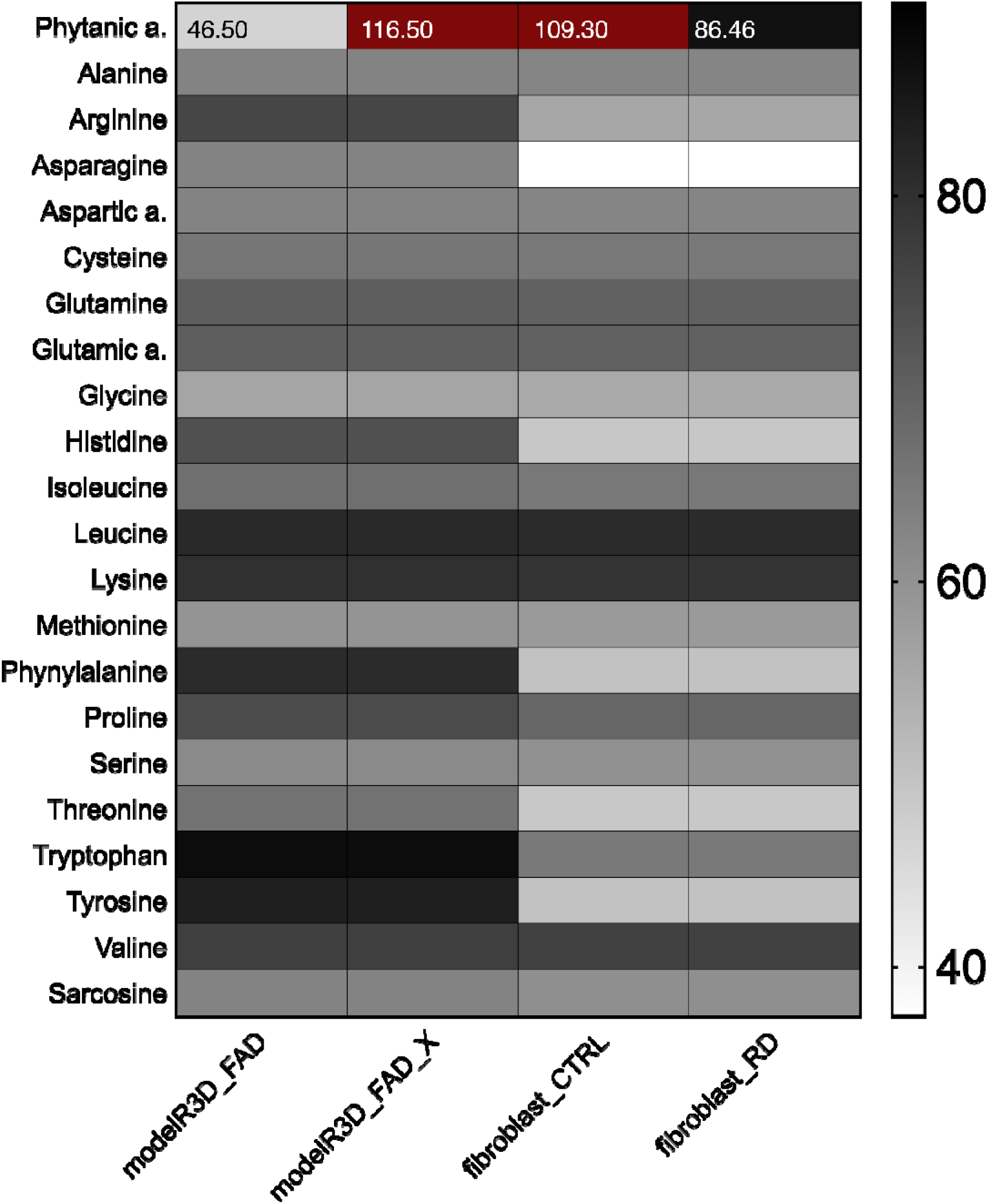
Model predictions of ATP yields from a single carbon source. Assessment of carbon source utilisation on minimal media supplemented with glutathione and pantothenic acid based on the ATP production from single-carbon source, including Recon3D_FAD, curated Recon3D_FAD for phytanic acid metabolism (Recon3D_FAD_x), the fibroblast-specific model for control (fibroblast_CTRL) and diseased conditions (fibroblast_RD). Grey shades in the heat-maps reflect the relative net ATP production ranging from no (white) to high (black), and very high ATP production (dark red).

**Figure S3.**
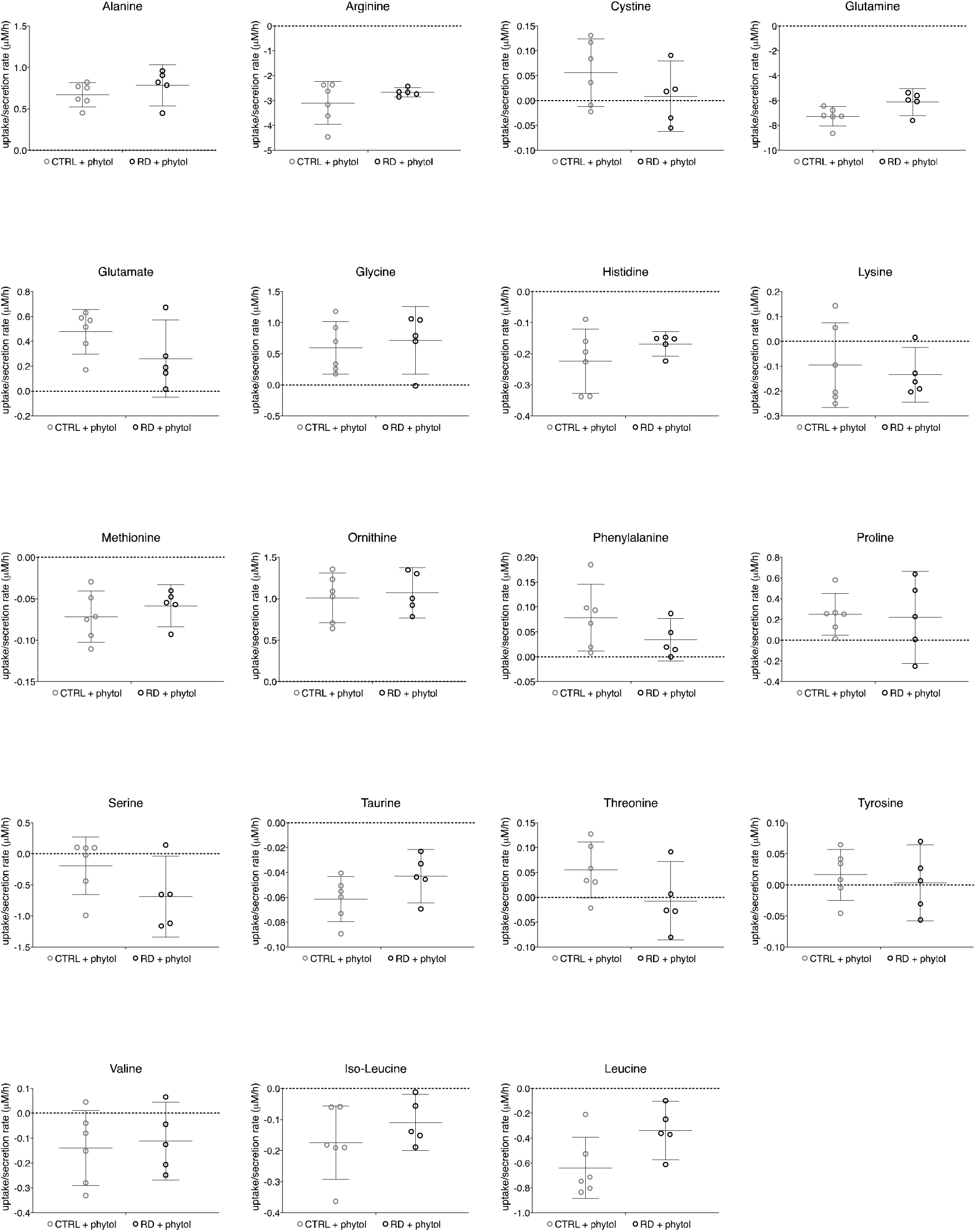
Additional data on amino acid uptake and secretion rates in the fibroblast CTRL and RD cultures exposed to phytol for 96 hours. Rates were calculated based on the fresh medium measurements.

**Figure S4.**
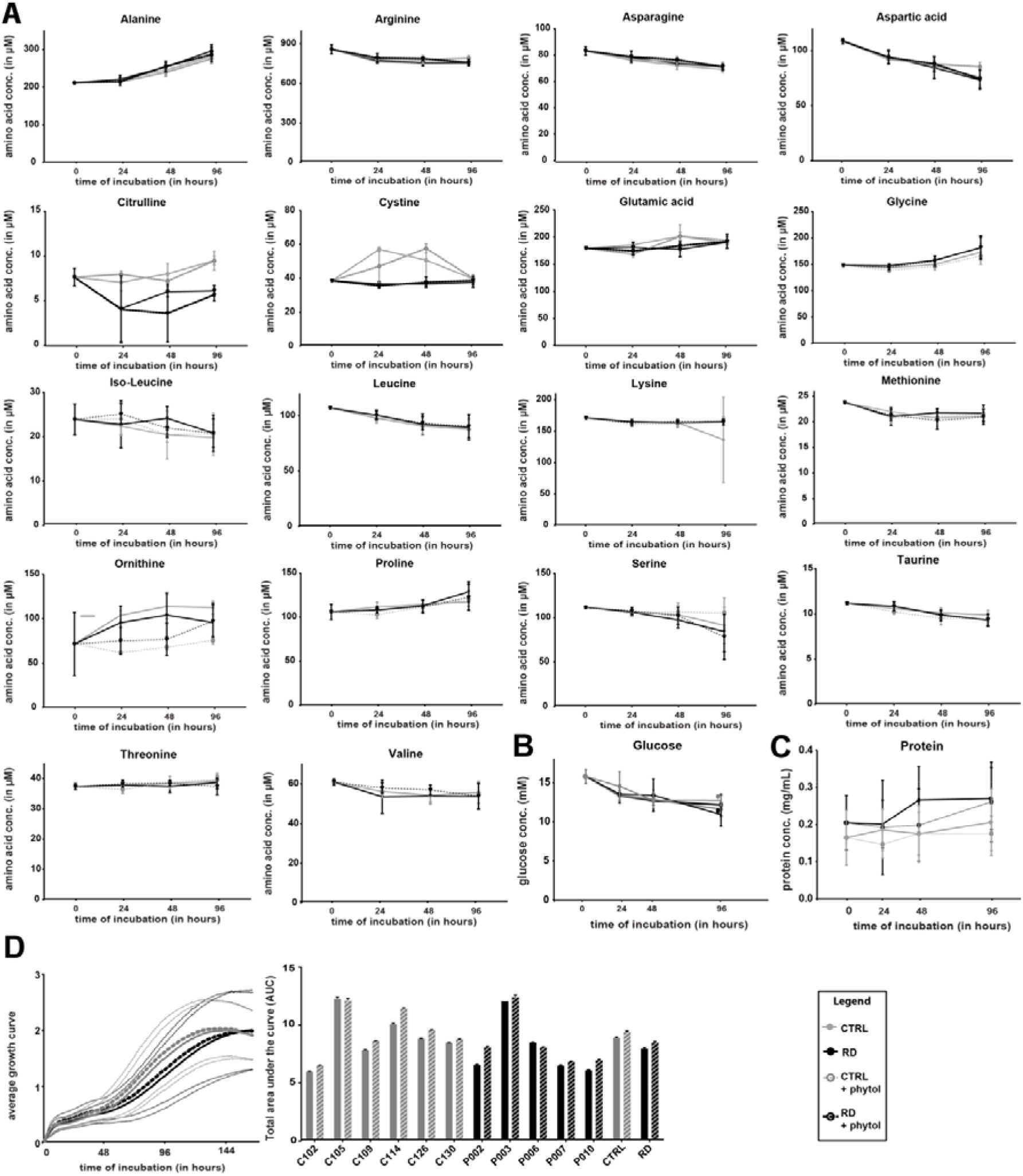
Additional experimental data of A) amino acids and B) glucose determinations in the medium of the cells after incubation at indicated time points. For details, see Fig. 3C. C) Protein concentrations of cell pellets after incubation at indicated time points. D) Growth curves of attached cells for the indicated time points (left panel), and statistical analysis of the total area under the curve per cell line after 7 days of incubation (right panel). Data are shown as bar plots.

## References

1. Wanders RJA, Ferdinandusse S, Brites P, Kemp S. Peroxisomes, lipid metabolism and lipotoxicity [Internet]. Biochimica et Biophysica Acta - Molecular and Cell Biology of Lipids. 2010. pp. 272–280. doi:10.1016/j.bbalip.2010.01.001

2. Van Veldhoven PP. Biochemistry and genetics of inherited disorders of peroxisomal fatty acid metabolism. J Lipid Res. 2010;51:2863–2895. doi:10.1194/jlr.R005959

3. Wanders RJA a, Komen J, Ferdinandusse S. Phytanic acid metabolism in health and disease. Biochim Biophys Acta. 2011;1811:498–507. doi:10.1016/j.bbalip.2011.06.006

4. Wanders RJA, Komen J, Kemp S. Fatty acid omega-oxidation as a rescue pathway for fatty acid oxidation disorders in humans. FEBS J. 2011;278:182–194. doi:10.1111/j.1742-4658.2010.07947.x

5. Wierzbicki ASA, Lloyd MDM, Schofield CJ, Feher MD, Gibberd FB. Refsum’s disease: a peroxisomal disorder affecting phytanic acid alpha-oxidation. J Neurochem. 2002;80:727–735. doi:10.1046/j.0022-3042.2002.00766.x

6. Baldwin EJ, Gibberd FB, Harley C, Sidey MC, Feher MD, Wierzbicki AS. The effectiveness of long-term dietary therapy in the treatment of adult Refsum disease. J Neurol Neurosurg Psychiatry. 2010;81:954–957. doi:10.1136/jnnp.2008.161059

7. Palsson BØ. Systems Biology: Properties of Reconstructed Networks. Cambridge University Press; 2006.

8. Thiele I, Swainston N, Fleming RMT, Hoppe A, Sahoo S, Aurich MK, et al. A community-driven global reconstruction of human metabolism. Nat Biotechnol. 2013;31:419–25. doi:10.1038/nbt.2488

9. Brunk E, Sahoo S, Zielinski DC, Altunkaya A, Dräger A, Mih N, et al. Recon3D enables a three-dimensional view of gene variation in human metabolism. Nat Biotechnol. 2018;36:272–281. doi: 10.1038/nbt.4072

10. Mardinoglu A, Agren R, Kampf C, Asplund A, Uhlen M, Nielsen J. Genome-scale metabolic modelling of hepatocytes reveals serine deficiency in patients with non-alcoholic fatty liver disease. Nat Commun. 2014;5:3083. doi:10.1038/ncomms4083

11. Edwards LM, Sigurdsson MI, Robbins PA, Weale ME, Cavalleri GL, Montgomery HE, et al. Genome-Scale Methods Converge on Key Mitochondrial Genes for the Survival of Human Cardiomyocytes in Hypoxia. Circ Cardiovasc Genet. 2014;7:407–415. doi: 10.1161/CIRCGENETICS.113.000269

12. Karlstädt A, Fliegner D, Kararigas G, Ruderisch HS, Regitz-Zagrosek V, Holzhütter H-G. CardioNet: a human metabolic network suited for the study of cardiomyocyte metabolism. BMC Syst Biol. 2012;6:114. doi:10.1186/1752-0509-6-114

13. Gille C, Bölling C, Hoppe A, Bulik S, Hoffmann S, Hübner K, et al. HepatoNet1: a comprehensive metabolic reconstruction of the human hepatocyte for the analysis of liver physiology. Mol Syst Biol. 2010;6:411. doi:10.1038/msb.2010.62

14. Sahoo S, Thiele I. Predicting the impact of diet and enzymopathies on human small intestinal epithelial cells. Hum Mol Genet. 2013;22:2705–2722. doi:10.1093/hmg/ddt119

15. Ryu JY, Kim HU, Lee SY. Reconstruction of genome-scale human metabolic models using omics data. Integr Biol. 2015;7:859–868. doi:10.1039/c5ib00002e

16. Dipple KM, McCabe ERB. Phenotypes of Patients with “Simple” Mendelian Disorders Are Complex Traits: Thresholds, Modifiers, and Systems Dynamics. Am J Hum Genet. 2000;66:1729–1735. doi:10.1086/302938

17. Sahoo S, Franzson L, Jonsson JJ, Thiele I, Sahoo S, Franzson L, et al. A compendium of inborn errors of metabolism mapped onto the human metabolic network. Mol Biosyst. 2012;8:2545. doi:10.1039/c2mb25075f

18. Argmann CA, Houten SM, Zhu J, Schadt EE. A Next Generation Multiscale View of Inborn Errors of Metabolism [Internet]. Cell Metabolism. NIH Public Access; 2016. pp. 13–26. doi:10.1016/j.cmet.2015.11.012

19. Shlomi T, Cabili MN, Ruppin E. Predicting metabolic biomarkers of human inborn errors of metabolism. Mol Syst Biol. 2009;5:263. doi:10.1038/msb.2009.22

20. Wegrzyn AB, Stolle S, Rienksma RA, Martins dos Santos VAP, Bakker BM, Suarez-Diez M. Cofactors revisited – Predicting the impact of flavoprotein-related diseases on a genome scale. Biochim Biophys Acta - Mol Basis Dis. 2019;1865. doi:10.1016/j.bbadis.2018.10.021

21. Pagliarini R, di Bernardo D. A Genome-Scale Modeling Approach to Study Inborn Errors of Liver Metabolism: Toward an In Silico Patient. J Comput Biol. 2013;20:383–397. doi:10.1089/cmb.2012.0276

22. Pagliarini R, Castello R, Napolitano F, Borzone R, Annunziata P, Mandrile G, et al. In Silico Modeling of Liver Metabolism in a Human Disease Reveals a Key Enzyme for Histidine and Histamine Homeostasis. Cell Rep. 2016;15:2292–2300. doi:10.1016/j.celrep.2016.05.014

23. Ferdinandusse S, Ebberink MS, Vaz FM, Waterham HR, Wanders RJA. The important role of biochemical and functional studies in the diagnostics of peroxisomal disorders. J Inherit Metab Dis. 2016;39:531–543. doi:10.1007/s10545-016-9922-4

24. van Roermund CWT, Ijlst L, Wagemans T, Wanders RJ a, Waterham HR. A role for the human peroxisomal half-transporter ABCD3 in the oxidation of dicarboxylic acids. Biochim Biophys Acta. 2014;1841:563–8. doi:10.1016/j.bbalip.2013.12.001

25. Ferdinandusse S, Denis S, Van Roermund CWT, Wanders RJA, Dacremont G. Identification of the peroxisomal beta-oxidation enzymes involved in the degradation of long-chain dicarboxylic acids. J Lipid Res. 2004;45:1104–11. doi:10.1194/jlr.M300512-JLR200

26. Wierzbicki AASAAS, Mayne PPDP, Lloyd MMDM, Burston D, Mei G, Sidey MC, et al. Metabolism of phytanic acid and 3-methyl-adipic acid excretion in patients with adult Refsum disease. J Lipid Res. 2003;44:1481–1488. doi:10.1194/jlr.M300121-JLR200

27. Uhlen M, Fagerberg L, Hallstrom BM, Lindskog C, Oksvold P, Mardinoglu A, et al. Tissue-based map of the human proteome. Science (80-). 2015;347:1260419–1260419. doi:10.1126/science.1260419

28. Matsumoto M, Matsuzaki F, Oshikawa K, Goshima N, Mori M, Kawamura Y, et al. A large-scale targeted proteomics assay resource based on an in vitro human proteome. Nat Meth. 2017;14:251–258. doi:10.1038/nmeth.4116

29. Amberger JS, Bocchini CA, Schiettecatte F, Scott AF, Hamosh A. OMIM.org: Online Mendelian Inheritance in Man (OMIM(®)), an online catalog of human genes and genetic disorders. Nucleic Acids Res. 2015;43:D789–D798. doi:10.1093/nar/gku1205

30. Vlassis N, Pacheco MP, Sauter T. Fast Reconstruction of Compact Context-Specific Metabolic Network Models. Ouzounis CA, editor. PLoS Comput Biol. 2014;10:e1003424. doi:10.1371/journal.pcbi.1003424

31. Lemons JMS, Feng X-JJ, Bennett BD, Legesse-Miller A, Johnson EL, Raitman I, et al. Quiescent Fibroblasts Exhibit High Metabolic Activity. PLOS Biol. 2010;8:e1000514. doi:10.1371/journal.pbio.1000514

32. Russell DW. Fifty years of advances in bile acid synthesis and metabolism. J Lipid Res. 2009;50 Suppl: S120–S125. doi:10.1194/jlr.R800026-JLR200

33. Kaufman DE, Smith RL. Direction choice for accelerated convergence in hit-and-run sampling. Oper Res. 1998;46:84–95. doi:10.1287/opre.46.1.84

34. Van Den Brink DM, Wanders RJA. Phytanic acid: Production from phytol, its breakdown and role in human disease. Cellular and Molecular Life Sciences. 2006. pp. 1752–1765. doi:10.1007/s00018-005-5463-y

35. Thiele I, Sahoo S, Heinken A, Heirendt L, Aurich MK, Noronha A, et al. When metabolism meets physiology: Harvey and Harvetta. bioRxiv. 2018; 255885. doi:10.1101/255885

36. Schönfeld P, Reiser G. Brain Lipotoxicity of Phytanic Acid and Very Long-chain Fatty Acids. Harmful Cellular/Mitochondrial Activities in Refsum Disease and X-Linked Adrenoleukodystrophy. Aging Dis. 2016;7:136. doi:10.14336/AD.2015.0823

37. Ferdinandusse S, Zomer AWM, Komen JC, van den Brink CE, Thanos M, Hamers FPT, et al. Ataxia with loss of Purkinje cells in a mouse model for Refsum disease. Proc Natl Acad Sci. 2008;105:17712–17717. doi:10.1073/pnas.0806066105

38. Swainston N, Smallbone K, Hefzi H, Dobson PD, Brewer J, Hanscho M, et al. Recon 2.2: from reconstruction to model of human metabolism. Metabolomics. 2016;12:109. doi:10.1007/s11306-016-1051-4

39. Sanders R-J, Ofman R, Dacremont G, Wanders RJA, Kemp S. Characterization of the human ω-oxidation pathway for ω-hydroxy-very-long-chain fatty acids. FASEB J. 2008;22:2064–2071. doi:10.1096/fj.07-099150

40. Sánchez A, Vázquez A. Bioactive peptides: A review. Food Qual Saf. 2017;1:29–46. doi:10.1093/fqsafe/fyx006

41. Weichart D, Gobom J, Klopfleisch S, Häsler R, Gustavsson N, Billmann S, et al. Analysis of NOD2-mediated proteome response to muramyl dipeptide in HEK293 cells. J Biol Chem. 2006;281:2380–9. doi:10.1074/jbc.M505986200

42. Bonfanti L, Peretto P, De Marchis S, Fasolo A. Carnosine-related dipeptides in the mammalian brain. Prog Neurobiol. 1999;59:333–53. Available: http://www.ncbi.nlm.nih.gov/pubmed/10501633

43. Snyder SH. Brain peptides as neurotransmitters. Science. 1980;209:976–83. doi:10.1126/science.6157191

44. Osborn MP, Park Y, Parks MB, Burgess LG, Uppal K, Lee K, et al. Metabolome-Wide Association Study of Neovascular Age-Related Macular Degeneration. Swaroop A, editor. PLoS One. 2013;8:e72737. doi:10.1371/journal.pone.0072737

45. Hušek P, Švagera Z, Všianský F, Franeková J, Šimek P. Prolyl-hydroxyproline dipeptide in non-hydrolyzed morning urine and its value in postmenopausal osteoporosis. Clin Chem Lab Med. 2008;46:1391–1397. doi:10.1515/CCLM.2008.259

46. Wu M, Xu Y, Fitch WL, Zheng M, Merritt RE, Shrager JB, et al. Liquid chromatography/mass spectrometry methods for measuring dipeptide abundance in non-small-cell lung cancer. Rapid Commun Mass Spectrom. 2013;27:2091–2098. doi:10.1002/rcm.6656

47. Fonteh AN, Harrington RJ, Tsai A, Liao P, Harrington MG. Free amino acid and dipeptide changes in the body fluids from Alzheimer’s disease subjects. Amino Acids. 2007;32:213–224. doi:10.1007/s00726-006-0409-8

48. Wierzbicki AS. Peroxisomal disorders affecting phytanic acid alpha-oxidation: a review. Biochem Soc Trans. 2007;35:881–886. doi:10.1042/BST0350881

49. Reiser G, Schönfeld P, Kahlert S. Mechanism of toxicity of the branched-chain fatty acid phytanic acid, a marker of Refsum disease, in astrocytes involves mitochondrial impairment. International journal of developmental …. 2006. pp. 113–22. doi:10.1016/j.ijdevneu.2005.11.002

50. Yates A, Akanni W, Amode MR, Barrell D, Billis K, Carvalho-Silva D, et al. Ensembl 2016. Nucleic Acids Res. 2016;44:D710–D716. doi:10.1093/nar/gkv1157

51. Dobin A, Davis CA, Schlesinger F, Drenkow J, Zaleski C, Jha S, et al. STAR: ultrafast universal RNA-seq aligner. Bioinformatics. 2013;29:15–21. doi:10.1093/bioinformatics/bts635

52. Li H, Handsaker B, Wysoker A, Fennell T, Ruan J, Homer N, et al. The Sequence Alignment/Map format and SAMtools. Bioinformatics. 2009;25:2078–2079. doi:10.1093/bioinformatics/btp352

53. Anders S, Pyl PT, Huber W. HTSeq-A Python framework to work with high-throughput sequencing data. Bioinformatics. 2015;31:166–169. doi:10.1093/bioinformatics/btu638

54. Andrews S. FastQC a Quality Control Tool for High Throughput Sequence Data [Internet]. 2010. Available: http://www.bioinformatics.babraham.ac.uk/projects/fastqc/

55. Broadinstitute. Pickard-tools [Internet]. 2016. Available: https://broadinstitute.github.io/picard/

56. Cox J, Mann M. MaxQuant enables high peptide identification rates, individualized p.p.b.- range mass accuracies and proteome-wide protein quantification. Nat Biotechnol. 2008;26:1367–72. doi:10.1038/nbt.1511

57. Cox J, Hein MY, Luber C a, Paron I, Nagaraj N, Mann M. Accurate Proteome-wide Label-free Quantification by Delayed Normalization and Maximal Peptide Ratio Extraction, Termed MaxLFQ. Mol Cell Proteomics. 2014;13:2513–2526. doi:10.1074/mcp.M113.031591

58. Love MI, Huber W, Anders S. Moderated estimation of fold change and dispersion for RNA-seq data with DESeq2. Genome Biol. 2014;15:550. doi:10.1186/s13059-014-0550-8

59. Ke N, Wang X, Xu X, Abassi YA. The xCELLigence System for Real-Time and Label-Free Monitoring of Cell Viability BT - Mammalian Cell Viability: Methods and Protocols. In: Stoddart MJ, editor. Totowa, NJ: Humana Press; 2011. pp. 33–43. doi:10.1007/978-1-61779-108-6_6

60. Moore S, Spackman DH, Stein WH. Chromatography of Amino Acids on Sulfonated Polystyrene Resins. An Improved System. Anal Chem. 1958;30:1185–1190. doi:10.1021/ac60139a005

61. Jansen G, Muskiet FAJ, Schierbeek H, Berger R, van der Slik W. Capillary gas chromatographic profiling of urinary, plasma and erythrocyte sugars and polyols as their trimethylsilyl derivatives, preceded by a simple and rapid prepurification method. Clin Chim Acta. 1986;157:277–293. doi:10.1016/0009-8981(86)90303-7

62. Vreken P, Van Lint AEM, Bootsma AH, Overmars H, Wanders RJA, Van Gennip AH. Rapid stable isotope dilution analysis of very-long-chain fatty acids, pristanic acid and phytanic acid using gas chromatography-electron impact mass spectrometry. J Chromatogr B Biomed Appl. 1998;713:281–287. doi:10.1016/S0378-4347(98)00186-8

62. Wanders RJA, Waterham HR, Ferdinandusse S. Metabolic Interplay between Peroxisomes and Other Subcellular Organelles Including Mitochondria and the Endoplasmic Reticulum. Front Cell Dev Biol. 2016;3: 83. doi:10.3389/fcell.2015.00083

64. Fleming RMT, Thiele I, Sahoo S, Haraldsd HS, Haraldsdóttir HS, Fleming RMT, et al. Modeling the effects of commonly used drugs on human metabolism. FEBS J. 2015;282:297–317. doi:10.1111/febs.13128

65. Bordbar A, Feist AM, Usaite-Black R, Woodcock J, Palsson BO, Famili I. A multi-tissue type genome-scale metabolic network for analysis of whole-body systems physiology. BMC Syst Biol. 2011;5:180. doi:10.1186/1752-0509-5-180

66. Uhlen M, Oksvold P, Fagerberg L, Lundberg E, Jonasson K, Forsberg M, et al. Towards a knowledge-based Human Protein Atlas. Nat Biotechnol. 2010;28:1248–1250. doi:10.1038/nbt1210-1248

67. The UniProt Consortium. UniProt: the universal protein knowledgebase. Nucleic Acids Res. 2017;45:D158–D169. Available: http://dx.doi.org/10.1093/nar/gkw1099

68. Heirendt L, Arreckx S, Pfau T, Mendoza SN, Richelle A, Heinken A, et al. Creation and analysis of biochemical constraint-based models using the COBRA Toolbox v.3.0. Nat Protoc. 2019;14:639–702. doi:10.1038/s41596-018-0098-2

69. Wanders RJA, van Roermund CWT, Schor DSM, ten Brink HJ, Jakobs C. 2-Hydroxyphytanic acid oxidase activity in rat and human liver and its deficiency in the Zellweger syndrome. Biochim Biophys Acta - Mol Basis Dis. 1994;1227:177–182. doi:10.1016/0925-4439(94)90092-2

70. Komen JC, Duran M, Wanders RJA. ω-Hydroxylation of phytanic acid in rat liver microsomes: implications for Refsum disease. J Lipid Res. 2004;45:1341–1346. doi:10.1194/jlr.M400064-JLR200

71. Perez-Riverol Y, Csordas A, Bai J, Bernal-Llinares M, Hewapathirana S, Kundu DJ, et al. The PRIDE database and related tools and resources in 2019: improving support for quantification data. Nucleic Acids Res. 2019;47:D442–D450. doi:10.1093/nar/gky1106

72. Barrett T, Wilhite SE, Ledoux P, Evangelista C, Kim IF, Tomashevsky M, et al. NCBI GEO: archive for functional genomics data sets--update. Nucleic Acids Res. 2013;41:D991–5. doi:10.1093/nar/gks1193

